# Mycobacterial infection induces a specific human innate immune response

**DOI:** 10.1101/017483

**Authors:** John D. Blischak, Ludovic Tailleux, Amy Mitrano, Luis B. Barreiro, Yoav Gilad

**Affiliations:** University of Chicago, Chicago, Illinois, 60637; Institut Pasteur, Mycobacterial Genetics Unit, rue du Dr Roux, F-75015 Paris, France; University of Montreal, Montreal, Québec, Canada

## Abstract

The innate immune system provides the first response to pathogen infection and orchestrates the activation of the adaptive immune system. Though a large component of the innate immune response is common to all infections, pathogen-specific responses have been documented as well. The innate immune response is thought to be especially critical for fighting infection with *Mycobacterium tuberculosis* (MTB), the causative agent of tuberculosis (TB). While TB can be deadly, only 5-10% of individuals infected with MTB develop active disease. The risk for disease susceptibility is, at least partly, heritable. Studies of inter-individual variation in the innate immune response to MTB infection may therefore shed light on the genetic basis for variation in susceptibility to TB. Yet, to date, we still do not know which properties of the innate immune response are specific to MTB infection and which represent a general response to pathogen infection. To begin addressing this gap, we infected macrophages with eight different bacteria, including different MTB strains and related mycobacteria, and studied the transcriptional response to infection. Although the ensued gene regulatory responses were largely consistent across the bacterial infection treatments, we were able to identify a novel subset of genes whose regulation was affected specifically by infection with mycobacteria. Genetic variants that are associated with regulatory differences in these genes should be considered candidate loci for explaining inter-individual susceptibility TB.

**Author Summary:** Tuberculosis (TB) is a deadly disease responsible for millions of deaths annually. It is caused by infection with *Mycobacterium tuberculosis* (MTB), an ancient human pathogen. Approximately a third of the world’s population is infected with MTB, yet only an estimated 5-10% of individuals will develop an active form of the disease. While this variation in TB susceptibility has been demonstrated to be heritable, we still know little about its underlying genetic basis. The genetic variation that affects TB susceptibility likely involves the innate immune system, which is our first line of defense against invading pathogens, because infection with MTB does not prevent future infections. However, we do not fully understand how the innate immune system differs in its response to MTB versus other bacteria. To investigate this further, we infected macrophages with MTB, related mycobacteria, and other bacteria, and measured how their gene expression levels changed in response. We identified a subset of genes that respond preferentially to infection with mycobacterial species. These genes provide insight into the interactions between MTB and the innate immune system and are candidate loci for explaining inter-individual susceptibility to TB.

## Introduction

The innate immune system provides the first line of defense against microbial pathogens. Broadly speaking, innate immune cells recognize foreign molecules through pattern recognition receptors (PRRs), e.g. Toll-like receptors (TLRs), which bind to highly-conserved pathogenic motifs known as pathogen-associated molecular patterns (PAMPs) [1,2]. In addition, innate immune cells recognize damage-associated molecular patterns (DAMPs) of host molecules released by infected cells [3]. The initial innate response involves the release of proinflammatory cytokines and lipids to recruit and activate other immune cells, phagocytosis of the pathogen, and apoptosis [4]. If the infection persists, the phagocytes stimulate the adaptive immune system by presenting antigens to activate T and B cells. In contrast to the highly specific adaptive immune response, the innate immune response has traditionally been viewed as a general response to infection.

Yet, more recent work revealed that the innate immune system also produces a pathogen-specific response in addition to the general response [5–8]. Furthermore, this pathogen-specific innate response can in turn affect the specificity of the adaptive immune response by directing the differentiation of T cells into distinct subtypes [9]. That said, though we developed an appreciation for the importance of the specific innate immune response, we still do not know the extent to which the innate immune response differs between infections nor fully understand the consequences of specific innate immune responses for fighting pathogens. One of the first challenges is to distinguish the unique immune response to a specific pathogen from the large core more general response.

The pathogen-specific innate immune response is determined, at least in part, by the specificity of the PRRs of the host immune cell. Each PRR binds to its specific targets and activates certain downstream signaling pathways [10]. For example, treatment of mouse dendritic cells with lipopolysaccharide (LPS), which is found on the outer membrane of gram-negative bacteria, or with PAM3CSK4 (PAM), which is a synthetic lipoprotein that mimics those found on both gram-negative and gram-positive bacteria, induce different transcriptional response programs, because the two antigens are bound by TLR4 and TLR2, respectively [11]. Different pathogens not only stimulate different PRRs, but they have also evolved different evasion mechanisms to manipulate the innate immune response [2,12–14]. These evasion strategies likely also contribute to the specificity of the response to different pathogens.

In the context of evasion strategies, the case of *Mycobacterium tuberculosis* (MTB), the causative agent of tuberculosis (TB), is especially interesting. In order to increase its success inside alveolar macrophages - the primary cells that target MTB upon infection - MTB prevents acidification of the macrophage phagosome by inhibiting vesicular proton pumps [12,15]. Additionally, MTB delays stimulation of the adaptive immune system by inducing host expression of anti-inflammatory cytokines [16,17]. Interestingly, while the adaptive immune system is needed to prevent the spread of MTB and subsequent onset of TB, infected individuals do not become immunized against future MTB infections. This property may be related to the difficulty to develop an effective vaccine for adult TB (the current vaccine, bacillus Calmette–Guérin, BCG, is partly effective in children, much less so in adults; Reference 18).

Interestingly from a human genetics viewpoint, there are large inter-individual differences in susceptibility to developing TB. While it is estimated that roughly a third of the human population is latently infected with MTB, only approximately 10% of healthy infected individuals will develop active TB (immunocompromised individuals, e.g. HIV-infected, develop active TB at a much higher frequency; Reference 19). Despite an inference for a strong individual genetic component to TB susceptibility, the genetic architecture remains largely unknown [20–23]. There have been quite a few reports of candidate-gene associations, but genome wide scans have only identified two weak associations with disease susceptibility [24–26].

To begin addressing this gap, we have previously investigated genetic variation that is associated with inter-individual differences in the transcriptional response of human phagocytes to infection with MTB [27]. We found 102 and 96 genes that were associated with an expression QTL (eQTL) only pre- or post-infection, respectively. We refer to these loci as response eQTLs since their association with gene expression is affected by MTB infection. Interestingly, these response eQTLs were enriched for significant signal in a genome wide association study of TB susceptibility [24]. However, it is unknown if the genes associated with these response eQTLs are induced specifically in response to infection with MTB or are a part of the core innate immune response.

In order to characterize the innate immune response specific to MTB infection and better understand the role of the response eQTL-associated genes in the innate immune response, we infected macrophages isolated from a panel of six healthy individuals with a variety of bacteria. In addition to MTB, we chose both related mycobacteria and more distantly related bacteria.

## Results

### Bacterial infection induces large changes in gene expression

To learn about the immune response to infection with different bacteria, with a particular emphasis on *Mycobacterium tuberculosis* (MTB), we investigated the *in vitro* gene regulatory response of macrophages to infection with multiple MTB stains, related mycobacterial species, and other bacterial species (Table 1). Specifically, we infected cultured macrophages with either MTB H37Rv, which is a common strain often used in laboratory experiments [28], MTB GC1237, which is a strain of the highly virulent Beijing family [29], bacillus Calmette-Guérin (BCG), which is attenuated *Mycobacterium bovis* used for vaccinations, *Mycobacterium smegmatis*, which is non-pathogenic, or heat-killed MTB H37Rv. In order to compare the response to infection with mycobacteria to the response to infection with other bacteria, we also included infection treatments with *Yersinia pseudotuberculosis* (gram-negative), *Salmonella typhimurium* (gram-negative), or *Staphylococcus epidermidis* (gram-positive).

**Table 1.**
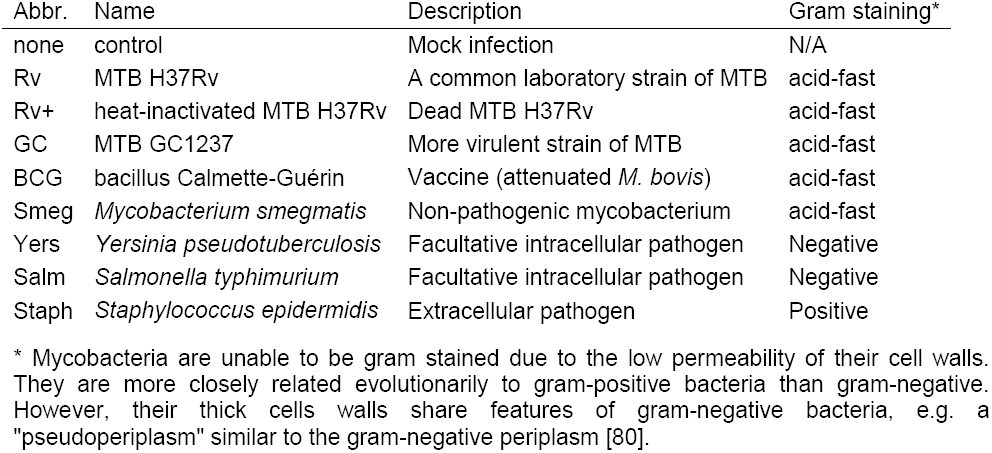
Description of bacteria.

We infected monocyte-derived macrophages from six individuals with the bacteria described above (including a non-infected control) and extracted RNA at 4, 18, and 48 hours post-infection (see Materials and Methods; Figure S1). We assessed RNA quality using the Agilent Bioanlyzer (Table S5) and sequenced the RNA to estimate gene expression levels. Detailed descriptions of our data processing, quality control analyses, and statistical modeling are available in the Materials and Methods section. Briefly, we mapped the short RNA-seq reads to the human genome (hg19) using the Subread algorithm [30], discarded reads that mapped non-uniquely, and counted the number of reads mapped to each protein-coding gene. We normalized the read counts using the weighted trimmed mean of M-values algorithm (TMM) [31], corrected for confounding “batch” effects, and used limma+voom [32–34] to test for differential expression (DE) between cultures infected with each bacteria and their time-matched controls (Table S2). Using this approach we initially observed the following general patterns: at four hours post-infection, only *Y. pseudotuberculosis* and *S. typhimurium* elicited a strong transcriptional response (Figure 1A); at 18 hours post-infection, all the bacteria had elicited a strong immune response (Figure 1A-B); and at 48 hours post-infection, all the bacteria continued stimulating the immune response (Figure 1A), however, many of the DE genes were not shared between the 18 and 48 hour timepoints (Figure 1C). Of note, at 48 hours post-infection we were unable to collect RNA from macrophages infected with *S. epidermidis* (see Materials and Methods).

**Figure 1.**
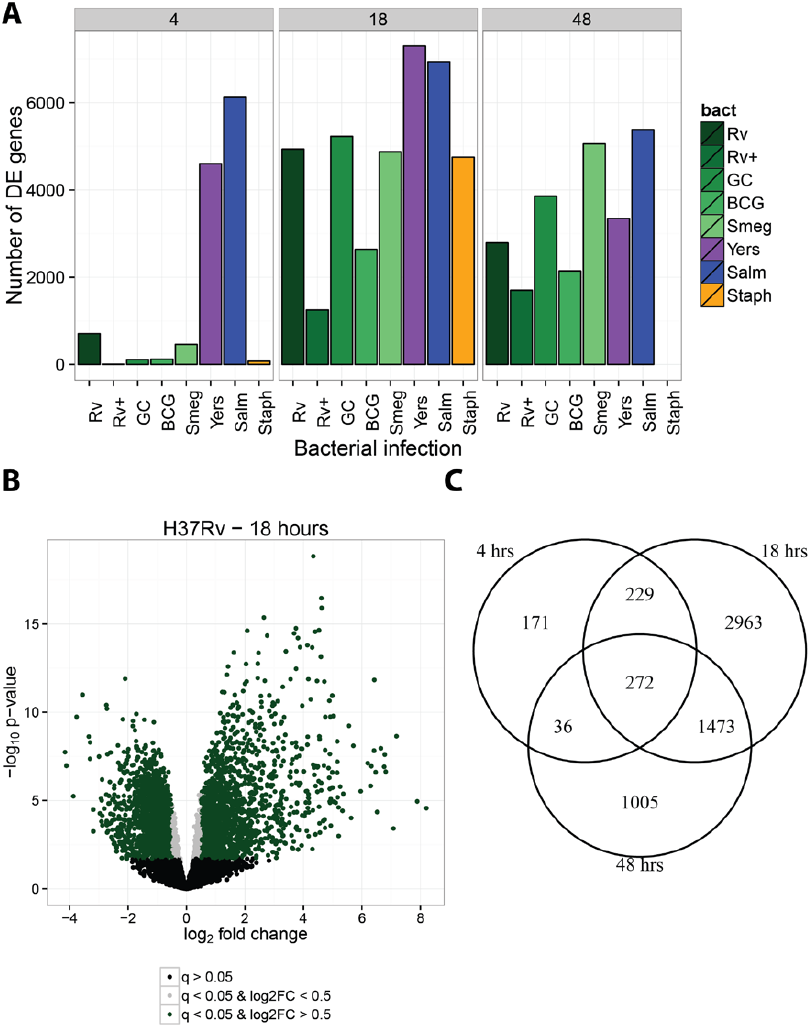
Differential expression analysis. We tested for differentially expressed genes for each bacterial infection by comparing it to its time-matched control. (A) We classified genes with q-value < 5% as differentially expressed. The mycobacteria are labeled in shades of green. (B) As expected, there were large transcriptional changes 18 hours post-infection with MTB H37Rv. Genes with q-value < 5% and absolute log_2_ fold change greater than 0.5 are labeled green, those with q-value < 5% and absolute log_2_ fold change less than 0.5 are labeled grey, and non-differentially expressed genes are labeled black. (C) The overlap in differentially expressed genes identified at 4, 18, and 48 hours post-infection with MTB H37Rv.

### Joint analysis identifies bacteria-specific response genes

In order to learn about variation in the innate immune response to bacterial infection, we identified genes whose regulation was altered by treatment with specific bacteria at specific timepoints. We first used a naïve approach whereby we determined all the pairwise overlaps between lists of DE genes across treatments (Table S6). The caveat of this strategy is that incomplete power can result in overestimating the difference between treatments. In order to account for incomplete power to detect DE genes when performing multiple pairwise comparisons, we performed a joint Bayesian analysis, which we implemented using the R/Bioconductor package Cormotif [35] (see Materials and Methods for more details). Using this approach, we classified genes into regulatory patterns based on their expression levels following each of the bacterial infections.

First, we examined the data across all the bacteria-time combinations. Initially we built a model that classified genes into one of 14 separate patterns based on their expression levels after each infection relative to their expression level in the non-infected control (Figure S2). However, we found that a model with only six expression patterns (Figure 2; Table S3), where a subset of the original 14 patterns are combined, is more intuitive from a biological perspective; thus we proceeded with the reduced model. Broadly speaking, we classified genes as responding in the early, middle, or late stages of infection, and we characterized the response as temporary or sustained. Pattern “non-DE” includes 4,245 genes whose expression levels were unchanged in all the experiments. Pattern “Yers-Salm” includes 1,414 early response genes whose expression levels changed at four hours post-infection with either *Y. pseudotuberculosis* or *S. typhimurium*, but not after infection with other bacteria. The genes in this pattern are enriched for gene ontology (GO) annotations related to type I interferon signaling (e.g. *SP100*, *IFI35*, *STAT2*), antigen presentation (*HLA-A*, *PSME1*, *CTSS*), and apoptosis (*CASP8*, *TRADD*, *FADD*) (Table S4). Pattern “18 h” includes 3,201 middle response genes whose expression levels changed exclusively at 18 hours post-infection in response to all bacteria and is enriched for GO annotations related to apoptosis (e.g. *E2F1*, *TP53*, *WWOX*). Pattern “48 h” includes 1,204 late response genes whose expression levels changed at 48 hours and is enriched for GO annotations related to phagocytosis (e.g. *MFGE8*, *COLEC12*) and tumor necrosis factor-mediated signaling (e.g. *STAT1*, *TRAF2*, *TNFRSF14*). Pattern “18 & 48 h” includes 1,926 middle-sustained response genes whose expression levels changed at 18 and 48 hours and is enriched for GO annotations related to the regulation of phagocytosis (e.g. *CD36*) and TLR signaling (*TLR1*, *TLR2*, *MYD88*). Lastly, pattern “All” includes 738 early-sustained genes whose expression levels changed after infection with all the bacteria across all three timepoints and is enriched for GO annotations related to type I interferon signaling (e.g. *IRF1*, *SOCS1*, *IFIT3*), cytokine secretion (*TNF*, *IL10*, *LILRB1*), and apoptosis (e.g. *IRF7*, *BCL2A1*, *MCL1*).

**Figure 2.**
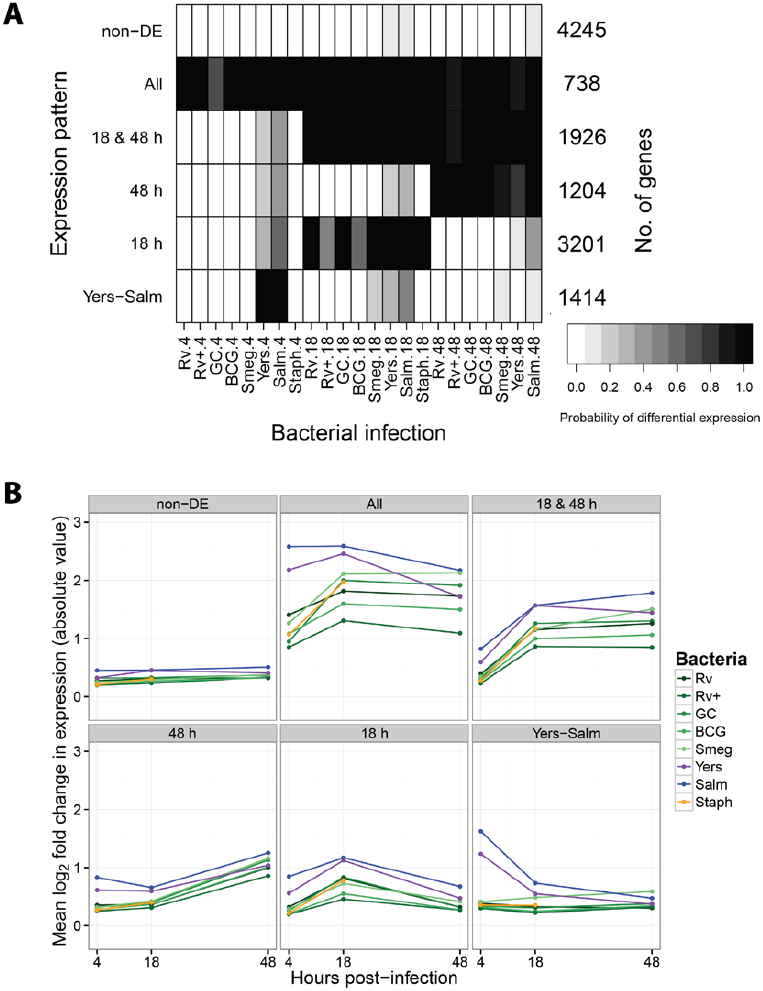
Joint Bayesian analysis. (A) Joint analysis of gene expression data from all three timepoints with Cormotif [35] identified six expression patterns: “non-DE”, “Yers-Salm”, “18 h”, “48 h”, “18 & 48 h”, and “All”. The shading of each box represents the posterior probability that a gene assigned to the expression pattern (row) is differentially expressed in response to infection with a particular bacteria (column), with black representing a high posterior probability and white a low posterior probability. (B) Each data point is the mean log_2_ fold change in expression (absolute value) in response to infection with the given bacteria for all the genes assigned to the particular expression pattern.

Next, we tested for more specific patterns by performing Cormotif separately on the data from the middle (18 h) and late (48 h) stages of infection. At 18 hours post-infection, we identified five separate expression patterns (Figure 3; Table S3). Pattern “non-DE” includes 5,268 genes whose expression levels were unchanged across all infections. Pattern “All” includes 4,424 genes whose expression levels were affected by all infections (e.g. *IL24*, *IRF2*, *TLR2*). Pattern “MTB” includes 177 genes whose expression levels changed specifically in response to infection with mycobacteria (e.g. *NCF2*, *TNFSF13*, *CSF1*). These genes had a high posterior probability of being DE 18 hours after infection with MTB H37Rv, heat-killed H37Rv, MTB GC1237, and BCG. Furthermore, the gray shading for *M. Smegmatis* (Figure 3A) signified an intermediate posterior probability for DE. In essence, this pattern is a merger of two sets of genes that were not large enough to be separated: one set that was DE across all five mycobacteria and another that was only DE after infection with the MTB strains and the closely-related BCG, but not *M. Smegmatis*. Pattern “Virulent”, in contrast, includes 1,165 genes whose expression levels were less strongly changed after infection with heat-inactivated MTB H37Rv or the attenuated vaccine strain BCG compared to the other bacteria (e.g. *IL1R1*, *IRF1*, *PILRB*). Also the genes in this category only have an intermediate probability of responding to the non-pathogenic *M. smegmatis*. Lastly, pattern “Yers-Salm” includes 1,694 genes whose expression levels changed preferentially after infection with *Y. pseudotuberculosis* or *S. typhimurium* (e.g. *TLR8*, *TGFB1*, *IL18*).

**Figure 3.**
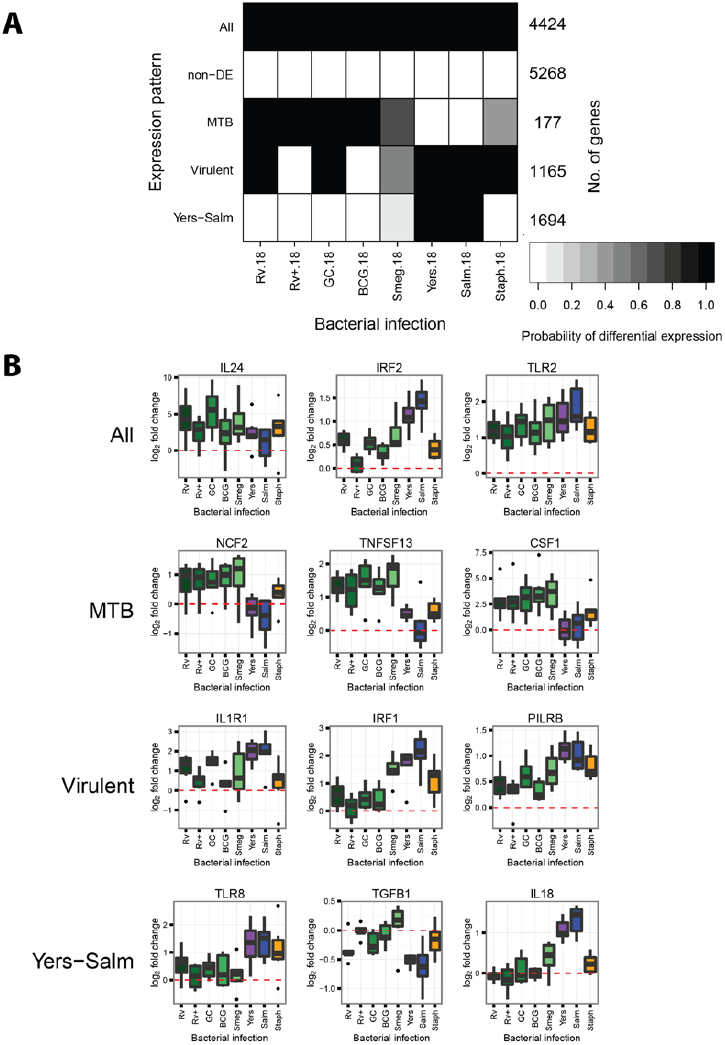
Joint Bayesian analysis - 18 hours post-infection. (A) Joint analysis of gene expression data from 18 hours post-infection with Cormotif identified five expression patterns: “Yers-Salm”, “Virulent”, “MTB”, “non-DE”, and “All”. (B) Example genes from the different expression patterns.

At 48 hours post-infection, we also discovered five expression patterns (Figure 4; Table S3). While many of the patterns have similar specificities to those observed at 18 hours post-infection, there is only little overlap across timepoints with respect to the genes comprising the patterns. For example, pattern “Yers-Salm” at 48 hours includes 1,582 genes whose expression levels changed strongly after infection with *Y. pseudotuberculosis* or *S. typhimurium* (e.g. *HLA-DPB1*, *IL10RB*, *CD248*), but only 263 of these genes are also in the corresponding pattern when we considered the data from the 18 hour timepoint. Similarly, at the 48 hour timepoint, pattern “MTB” includes 288 genes whose expression levels changed preferentially after infection with mycobacteria (e.g. *CCL1*, *ATP6V1A*, *IL27RA*), but only 33 of these genes are in the corresponding pattern at the 18 hour timepoint. Pattern “Virulent” includes 14 genes whose expression levels were not changed after infection with heat-inactivated MTB H37Rv or the attenuated vaccine strain BCG (e.g. *MAP3K4*, *SEMA4G*, *BTG1*), and only one of these also belongs to the pattern “Virulent” at 18 hours post-infection.

**Figure 4.**
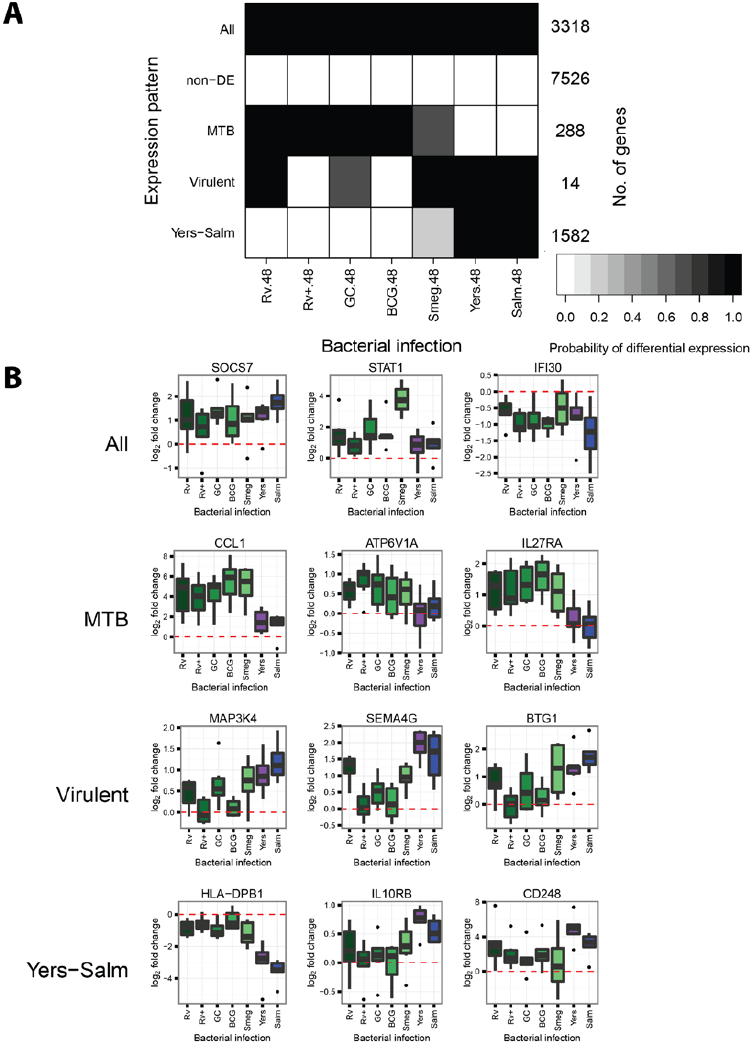
Joint Bayesian analysis - 48 hours post-infection. (A) Joint analysis of gene expression data from 48 hours post-infection with Cormotif identified five expression patterns: “Yers-Salm”, “Virulent”, “MTB”, “non-DE”, and “All”. (B) Example genes from the different expression patterns.

### Infection-induced response eQTLs are shared across bacterial infections

Using the gene expression patterns we identified by applying the joint analysis approach, we investigated the specificity of previously identified response eQTLs to infection with MTB H37Rv [27]. Since the response eQTLs were identified at 18 hours post-infection, we investigated the distribution of genes associated with response eQTLs among the five patterns we found at that timepoint (Figure 5A). Only one gene associated with a response eQTL was also DE specifically in response to MTB (*CMAS*). Otherwise, most of the response eQTL-associated genes were classified as either DE following infection with all bacteria or not DE in any infection. That a large proportion of the genes associated with response eQTLs were not DE in any of these experiments is likely due to the fact that the eQTL study was performed in dendritic cells whereas our data were collected from macrophages. Overall, our observations suggest that most of the previously identified response eQTLs are genetic variants that affect the human innate immune response to bacterial infection in general, and not specifically the response to MTB H37Rv.

**Figure 5.**
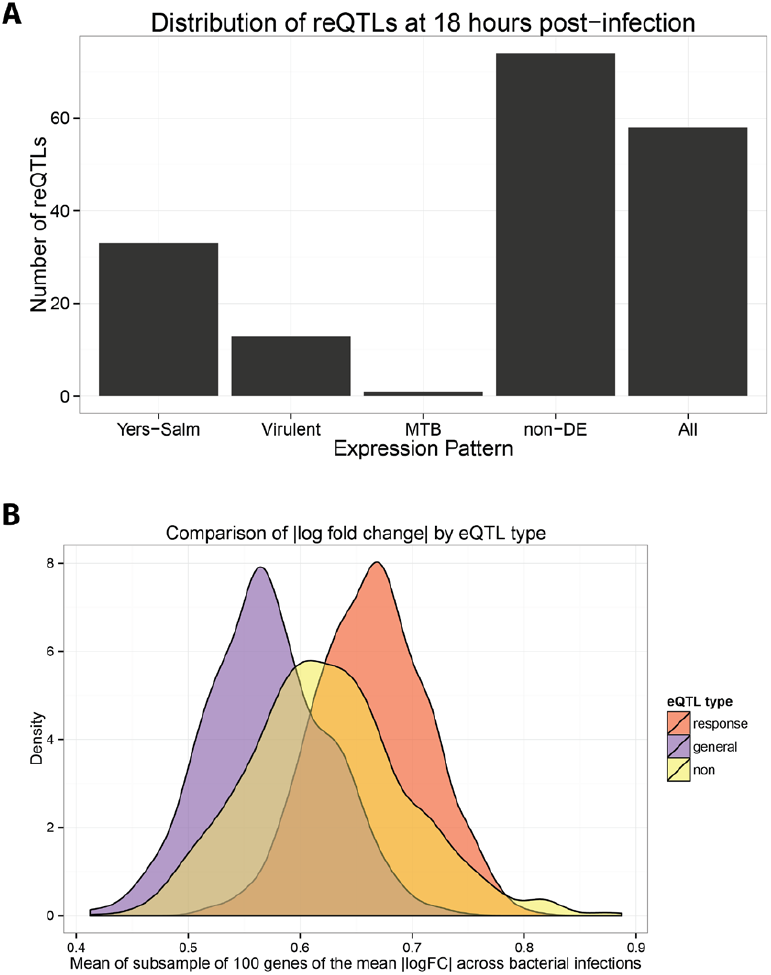
Response eQTLs at 18 hours post-infection. (A) We counted the number of response eQTLs from Barreiro et al. [27] (179 out of the 198 were also expressed in our study) in each of the five gene expression patterns at 18 hours post-infection (Figure 3). (B) We compared the mean log_2_ fold change in expression across the 8 bacterial infections at the 18 hour timepoint for three classes of genes: response eQTL genes (red), general eQTL genes (purple), and non-eQTL genes (yellow) (see Materials and Methods for details).

To provide further broad support for the interpretation that the response eQTL genes are important for the innate response to bacterial infection in general, we considered the log_2_ fold change in expression values following infection (Figure 1). For each gene, we calculated the mean log_2_ fold change in expression level across the eight bacterial infections at the 18 hour timepoint. Next, we compared the absolute values of the log_2_ fold change in expression between genes associated with response eQTLs, genes associated with general eQTLs (i.e. genes associated with an eQTL pre- and post-infection), and genes not found to be associated with an eQTL. Since there was a large difference in the number of genes in these three classes, we subsampled genes from each and calculated the mean of the absolute values (and repeated this process 1000 times). We found that the expression level of genes associated with response eQTLs is altered to a larger degree (significantly higher effect size; *P* < 2.2 × e^-16^; Figure 5B) following infection compared to the genes in the other two classes.

## Discussion

### Bayesian analysis identified mycobacteria-specific response genes

In order to identify general and treatment-specific gene regulatory responses, we performed a joint Bayesian analysis of the data using Cormotif [35]. By jointly analyzing the data, as opposed to comparing overlaps between independent lists of differentially expressed genes generated using an arbitrary cutoff, we minimized the identification of specific responses due to false negatives (i.e. genes that appear to be differentially expressed in response to a subset of bacterial infections when in reality the response is similar across all the infections). Similar to previous observations [5,6], we found a large core transcriptional response to infection. However, we also identified a novel subset of genes whose regulation is preferentially altered in response to infection with mycobacteria but not to the other bacteria we tested. Since these responses are unique to infection with mycobacteria (at least in the context of our study design), they may be promising candidates for future studies that focus on the mechanisms by which mycobacteria successfully subvert the human innate immune response.

One of these potential candidate genes caught our attention in particular. The neutrophil cytosolic factor 2 (*NCF2*, also known as *p67phox*) gene, whose expression levels were affected specifically by infection with mycobacteria at 18 hours post-infection, is a subunit of the phagocyte NAPDH oxidase. The phagocyte NAPDH oxidase is responsible for generating reactive oxygen species, which are utilized to fight intracellular pathogens [36–41]. These reactive oxygen species may also serve a signaling role in activating other immune cell types to ensure proper granuloma formation and killing of mycobacteria [42]. Loss-of-function mutations in subunits of the NAPDH oxidase cause chronic granulomatous disease (CGD) [41], which is characterized by the formation of granulomas throughout the body due to the inability of phagocytes to kill the ingested pathogens. In contrast to wild type animals, mice with mutations in subunits of the phagocyte NAPDH oxidase develop tuberculosis after infection with the vaccine strain, BCG [42]. Humans who are administered the vaccine before being diagnosed with CGD also develop the disease [41].

Another gene of interest is chemokine (C-C motif) ligand 1 (*CCL1*), which stimulates migration of human monocytes [43]. The expression of *CCL1* was increased in response to infection with any of the bacteria at 4 and 18 hours post-infection. However, at 48 hours post-infection, *CCL1* was assigned to the expression pattern “MTB” because its expression remained elevated in cells infected with one of the mycobacteria but not in cells infected with *S. typhimurium* or *Y. pseudotuberculosis* (Figure 4B). Interestingly, Thuong et al. identified *CCL1* as being induced to a greater exent in MTB-infected macrophages (4 hours post-infection) isolated from individuals with pulmonary TB compared to macrophages from individuals with latent TB infections [44]. Put together, our observations and those of Thuong and colleagues suggest that *CCL1* is involved in the pathogenesis of TB. Further supporting this notion, Thuong et al. also found a genetic association between variants in the *CCL1* region and TB susceptibility [44]. However to date, subsequent genetic association studies investigating *CCL1* have reported mixed results [45,46].

One caveat of the joint Bayesian analysis is that we were not able to classify genes into unusual patterns (because this approach can only discover expression patterns shared by a large number of genes and, by definition, only few genes fall into “unusual” patterns). For example, unusual patterns of interest include changes in expression specifically in response to some but not all of the mycobacterial infections. One gene that satisfied this pattern is the dual specificity phosphatase 14 (*DUSP14*). We specifically examined the expression data for this gene because it was previously associated with an MTB infection response eQTL in dendritic cells [27], and consequently when the eQTL results were used as a prior - *DUSP14* was found to be significantly associated with TB susceptibility. Moreover, knocking down *DUSP14* expression via siRNA in murine macrophages resulted in a lower bacterial load 90 hours post-infection with MTB H37Rv [47]. In our joint Bayesian analysis, *DUSP14* was not classified as one of the genes whose regulation was altered in response to infection with mycobacteria. Yet, *DUSP14* was upregulated at 18 hours post-infection with MTB H37Rv (q-value: 16%), MTB GC1237 (q-value: 3%), and BCG (q-value: 9%); and downregulated post-infection with *S. typhimurium* (q-value: 9%) (Figure S3). Thus, our data lends further support for the role of *DUSP14* as a TB susceptibility gene.

### Little evidence for strain-specific transcriptional response to infection

There are six major families of MTB that differ in their geographic distribution and virulence [48,49]. Strains from these families are known to differ in their growth rates inside macrophages [50], expression levels of bacterial genes [51,52], and cell wall lipid composition [53]. Previous studies have found that different MTB strains induce different innate immune responses in human cell lines and other infection models [54]. A dominate narrative is that MTB strains from East Asia, referred to as the Beijing family (Gagneux et al. classified it as MTB lineage 2 [48]), are more virulent because they induce a lower proinflammatory immune response compared to the common laboratory strains [55–59]. However, other studies have reported the opposite, namely that Beijing strains induce a larger proinflammatory response [60], or a conflicting response in which various pro- and anti-inflammatory cytokines are differentially regulated [61,62] compared to laboratory strains.

In our study, albeit with a small sample size, we found no marked differences between the transcriptional response to infection with MTB H37Rv or MTB GC1237, a Beijing strain (Figure S4; Table S7). Furthermore, the pro-inflammatory cytokines *TNF-α* and *IL-6* and the antiinflammatory cytokine *IL-10* were strongly upregulated in response to both strains of MTB (Figure S5). This observation is in concordance with Wu et al., who also reported no apparent difference in the transcriptional response of THP-1 cells to infection with MTB H37Rv versus multiple Beijing strains [63]. Thus the increased virulence of the Beijing family of MTB strains may be due to mechanisms not assayed in this study such as post-transcriptional effects, cell-cell signaling, and environmental stimuli. It should be noted, however, that not all Beijing strains are equally virulent [64,65] and that MTB H37Rv is a laboratory-adapted strain that has evolved independently in different laboratories [66].

### Previously identified response eQTLs affect response to bacterial infection in general

In a previous study, we identified response eQTLs that were associated with gene expression levels in MTB-infected human dendritic cells. We investigated the expression pattern of genes associated with the response eQTLs in our study. Using the five expression patterns identified by the joint Bayesian analysis at 18 hours post-infection, we examined the distribution of response eQTL genes and discovered that these genes were not enriched in the mycobacteria-specific expression pattern (Figure 5A). Instead, many were differentially expressed across all the infections (pattern “All”). Thus, response eQTLs modulate the inter-individual response to infection with diverse types of bacteria. That said, one gene was both associated with a response eQTL and specifically differentially expressed following mycobacterial infection. Though this result does not represent a significant enrichment of response eQTL genes among those whose regulation was affected specifically by infection with MTB, the identity of the gene renders the observation intriguing. *CMAS* (cytidine monophosphate N-acetylneuraminic acid synthetase), is an enzyme that is involved in the processing of sialic acid, which is then added to cell surface glycoproteins and glycolipids. Glycoproteins are known to be important in many functions of the immune response, including initial pathogen detection (e.g. TLRs) and antigen presentation (e.g. major histocompatibility complex (MHC) molecules) [67–69]. We suggest that this gene is an interesting candidate for further understanding both MTB pathogenesis and inter-individual susceptibility to tuberculosis.

## Materials and Methods

### Ethics Statement

Buffy coats were obtained from healthy donors after informed consent (Etablissement Français du Sang). In conformity with French regulations, the biobank has been declared to and recorded by both the French Ministry of Research and a French Ethics Committee under the reference DC-2008-68 collection 2.

### Sample collection and macrophage differentiation

We collected buffy coats (∼50 mL) from six healthy donors. Next we isolated peripheral blood mononuclear cells (PBMCs) via Ficoll-Paque centrifugation [28] and enriched for monocytes via positive selection with beads containing CD14 antibodies [27]. Then we differentiated the monocytes into macrophages by culturing for 6-7 days in RPMI buffer supplemented with macrophage colony-stimulating factor (M-CSF) [70].

### Bacterial infection

For each bacterial infection (Table 1), we treated the macrophages with a multiplicity of infection (MOI) of 2:1. After one hour, we washed the macrophages five times with phosphate-buffered saline (PBS) and treated them with gentamycin (50 μg/μL) to kill all extracellular bacteria. After one hour of antibiotic treatment, we changed the medium to a lower concentration of gentamycin (5 μg/μL), which marked the zero timepoint of the study. We allowed the cells to grow for 4, 18, or 48 hours before lysing them with QIAzol Lysis Reagent and then storing them at -80° C. We chose these timepoints based on a previous analysis of the human transcriptional response to infection with MTB [71]. No data is available for 48 hours post-infection with *S. epidermidis*. After escaping the macrophages upon cell death, sufficient *S. epidermidis* were able to proliferate in the gentmycin-supplemented medium to contiminate the entire well by 48 hours post-infection.

### RNA extraction, library preparation, and sequencing

We extracted RNA using the QIAgen miRNeasy kit. There were a total of 13 batches of 12 samples each (6 individuals × 9 conditions × 3 timepoints, minus 48 hours post-infection with *S. epidermidis*). We designed the batches to maximally partition the variables of interest (individual, condition, timepoint) in order to minimize the introduction of biases due to batch processing [72]. To assess RNA quality, we measured the RNA Integrity Number (RIN) with the Agilent Bioanalyzer (Table S5). Importantly, there were no significant differences in the RIN (mean of 7.8 ± 2.0) between the bacterial infections or between the timepoints (Figure S6B). In batches of 12 samples, we added barcoded adapters (Illumina TruSeq RNA Sample Preparation Kit v2) and sequenced 50 base pairs single end over multiple flow cells on the Illumina HiSeq 2500.

### Mapping, counting, and normalization

We mapped the short reads to the human genome (hg19) using the Subread algorithm [30] and discarded those that mapped non-uniquely. Next, we obtained the read counts for each Ensembl protein-coding gene (biotype: “protein_coding”) with the featureCounts algorithm, which sums the reads falling in the union of all exons of a gene and discards reads mapping to more than one gene [73]. There were no significant differences in the number of mapped exonic reads (mean of 41.8 ± 21.2 million per sample) between the bacterial infections or between the timepoints (Figure S6A). We removed genes with fewer than one count per million exonic reads in fewer than six samples. To account for differences in the read counts at the extremes of the distribution, we normalized the samples using the weighted trimmed mean of M-values algorithm (TMM) [31].

### Differential expression analysis

To assess the quality of the data, we performed principal components analysis (PCA) of the TMM-normalized log_2_-transformed counts per million (CPM). PC2 separated the samples by timepoint, but PC1 was associated with the RIN score and the processing batch (Figure S7A). After the effect of RIN score and processing batch was removed with the function removeBatchEffect from the limma package [74], PC1 separated the samples by timepoint and PC2 separated the infected and control samples (Figure S7B). All figures displaying expression data were generated using the batch-corrected data.

For the standard analysis, we tested for differential expression using limma+voom [32–34] because it has been shown to perform well with sufficient sample size (n >= 3 per condition) [75,76]. Based on the PCA results, we included RIN score and processing batch as covariates in the model. We corrected for multiple testing with the Benjamini & Hochberg false discovery rate (FDR) [77] and considered genes with q-value less than 5% to be differentially expressed.

Since we were interested in the shared and differential response to infection with the different bacteria, we performed a joint Bayesian analysis using the Cormotif algorithm [35]. Cormotif shares information across experiments, in this case infections, to identify the main patterns of differential gene expression (which it refers to as *correlation motifs*) and assigns each gene to one of these gene expression patterns. One caveat of the Cormotif algorithm is that is does not distinguish the direction of the effect across infections. In other words, a gene that is assigned to an expression pattern could be differentially expressed in different directions across the infections. However, in this data set, this was rarely observed (Table S8).

In practice, we had to make several modifications when using Cormotif. First, since the method was developed for microarray data, we used the batch-corrected TMM-normalized log_2_-transformed CPM as input. Second, the method assumes independence between the experiments, and we only have one control per timepoint. However, since this dependence will cause genes to be more likely to be either uniformly differentially expressed across all the infections or uniformly unchanged, this caveat is conservative to our results of gene expression patterns that are specific to subgroups of the bacterial infections. Third, the current version of the method (v1.12.0) does not return the cluster likelihoods, i.e. the likelihood that a gene belongs to each of the gene expression patterns. To facilitate downstream analyses with these sets of genes, we modified the original code to additionally return this information. Lastly, Cormotif is non-deterministic. Thus to obtain consistent results, we ran each test 100 times and kept the result with the largest maximum likelihood estimate.

We tested for enrichment of gene ontology (GO) biological processes among the genes in the gene expression patterns using topGO [78]. We tested for significance with the Fisher’s Exact Test, used the weight01 algorithm from topGO to account for the correlation among GO categories due to its graph structure, and considered significant any category with p-value less than 0.01.

### Analysis using previously identified response eQTLs

We downloaded the list of response eQTL genes from Supplementary Table 3 from Barreiro et al. [27]. Of the 198 response eQTL genes discovered in the dendritic cells in that study, 179 of the genes were also expressed in the macrophages from this study. In order to compare the differential expression results of the response eQTL genes to other genes, we used the log_2_ fold changes in expression estimated by limma [74]. First, we calculated the mean log_2_ fold change at 18 hours post-infection for each gene across the eight bacteria. Second, we converted these mean estimates to their absolute values. Third, we subsampled 100 genes from each of the three categories (response eQTL, general eQTL, and non-eQTL genes) and calculated the mean of the absolute values. We performed this subsampling 1000 times (Figure 5B). Fourth, we performed t-tests to compare the distribution of response eQTL genes to either that of the general eQTL genes or the non-eQTL genes.

### Data and code availability

The data have been deposited in NCBI’s Gene Expression Omnibus [79] and are accessible through GEO Series accession number GSE67427 (http://www.ncbi.nlm.nih.gov/geo/query/acc.cgi?acc=GSE67427). The code is available at https://bitbucket.org/jdblischak/tb.

## Author Contributions

YG, LT, and LBB conceived of the study and designed the experiments. LT performed the infection experiments. JDB extracted the RNA and analyzed the data. AM prepared the sequencing libraries. LBB and YG supervised the project. JDB and YG wrote the paper with input from all authors.

## Acknowledgments

We thank Matthew Stephens and Bryce van de Geijn for advice on the statistical analyses, and all members of the Gilad lab for helpful discussions. This work was supported by grant AI087658 to YG and LT and LBB. JDB was partially supported by National Institutes of Health Grant T32 GM007197.

